# Discovery of isoplumbagin as a novel NQO1 substrate and anti-cancer quinone

**DOI:** 10.1101/2020.04.28.063222

**Authors:** Yen-Chi Tsao, Yu-Jung Chang, Chun-Hsien Wang, Linyi Chen

## Abstract

Isoplumbagin (5-hydroxy-3-methyl-1,4-naphthoquinone), a naturally occurring quinone from *Lawsonia inermis* and *Plumbago europaea*, that has been reported to have anti-inflammatory and anti-microbial activity. Inflammation has long been implicated in cancer progression. In this study, we examined the anti-cancer effect of chemically-synthesized isoplumbagin. Our results revealed that isoplumbagin treatment suppressed cell viability and invasion of highly invasive oral squamous cell carcinoma (OSCC) OC3-IV2 cells, glioblastoma U87 cells, non-small cell lung carcinoma H1299 cells, prostate cancer PC3 cells, and cervical cancer Hela cells by using MTT and Boyden chamber assays. *In vivo* studies demonstrate the inhibitory effect of 2 mg/kg isoplumbagin on the growth of orthotopic xenograft tumors derived from OSCC cells. Mechanistically, isoplumbagin exerts its cytotoxic effect through acting as a substrate of NAD(P)H quinone dehydrogenase 1 (NQO1) to generate hydroquinone, which reverses mitochondrial fission phenotype, reduces mitochondrial complex IV activity and thus compromises mitochondrial function. Collectively, this work reveals an anti-cancer activity of isoplumbagin mainly through modulating mitochondrial dynamics and function.

**Chemical compounds:** Isoplumbagin (PubChem CID: 375105)

## 1. Introduction

Medicinal plants and their metabolites are great sources for pharmaceutical applications. The metabolites in plants provide rich variety of bioactive compounds with anti-cancer, anti-oxidant, anti-inflammatory or anti-microbial activities. Some of these drugs such as paclitaxel, docetaxel, vincristine, and vinblastine are approved and used extensively in treating several types of cancer, including breast, testicular and bladder cancers [1]. Natural quinones are secondary metabolites of plant and are categorized as benzoquinone, naphthoquinone, phenanthrenequinone, and anthraquinone according to their aromatic carbon skeleton [2]. Quinones are highly electrophilic molecules that accept one- or two-electron from flavoenzymes and iron-sulfur proteins to form semiquinone or hydroquinone. They exert cytotoxic effects through alkylating proteins or DNA, and affect redox cycle with their semiquinone radicals to generate reactive oxygen species. Their cytotoxicity promotes inflammatory reactions, oxidizes DNA, and induces cell death. Quinone-based drugs such as doxorubicin and mitomycin C have been used clinically for cancer chemotherapy, but their adverse side effects and toxicity have been an issue [3]. Consequently, there is a continued search for development of quinone-based agents displaying anti-tumor activity that are less toxic and reduced side effects.

Isoplumbagin (5-hydroxy-3-methyl-1,4-naphthoquinone) can be isolated from the bark of *Lawsonia inermis* [4] and *Plumbago europaea* [5] and has been shown to exhibit anti-inflammatory activity against Carrageenan-induced rat paw oedema [6] and anti-microbial activity against invasive vaginitis strains [5]. Chronic inflammation facilitates the initiation and progression of cancer [7]. Although isoplumbagin is known to exert pharmaceutical effects, no report thus far demonstrates its anti-cancer effect. Therefore, this study aims to evaluate its potential anti-cancer effect and the underlying mechanism.

## 2. Materials and methods

### 2.1. Antibodies and reagents

Anti-TOM20 (#sc-11415, 1:200 for immunofluorescence staining) and anti-NQO1 (#sc-32793, 1:1000 for immunoblotting) were purchased from Santa Cruz Biotechnology (Santa Cruz, CA, USA). Anti-GAPDH (#10494-1-AP, 1:20000 for immunoblotting) was purchased from Proteintech (Rosemont, IL, USA). Alexa Fluor 488 (#A11001) or Alexa Fluor 700 (#A21036)–conjugated secondary antibodies, Alexa Fluor 488–conjugated phalloidin (#A12379) and 4’,6-diamidino-2-phenylindole (DAPI) (#1306) were obtained from Invitrogen (Carlsbad, CA, USA). IRDye800CW-labeled anti-rabbit secondary antibody (#926-32211) was purchased from LI-COR Biosciences (Lincoln, NE, USA). Isoplumbagin was acquired from AKos GmbH (Steinen, Germany). Oligomycin (#75351), carbonyl cyanide 4-(trifluoromethoxy)phenylhydrazone (FCCP) (#C2920), rotenone (#R8875), antimycin A (#A8674), lactobionic acid (#153516), taurine (#T8691), digitonin (#D141), glutamate (#G8415), malate (#M1000), succinate (#S3674), adenosine 5’-diphosphate sodium salt (ADP) (#A2754), TMPD (#T3134) and ascorbate (#A4034) were purchased from Sigma-Aldrich (St Louis, MO, USA).

### 2.2. Cell culture

Human oral cancer cell line OC3-IV2 cells have been described [8–10] and were cultured in a 1:1 ratio of Dulbecco’s modified Eagle medium (DMEM; Invitrogen) and keratinocyte serum-free medium (KSFM; Invitrogen) containing 10% (v/v) fetal bovine serum (Invitrogen), 1% (v/v) L-glutamine (Invitrogen), and 1% (v/v) antibiotic-antimycotic (Invitrogen). Human primary glioblastoma cell line U87 were obtained from the American Type Culture Collection (ATCC) and grown in DMEM medium supplemented with 10% (v/v) fetal bovine serum, 1% (v/v) MEM Non-Essential Amino Acids Solution (Invitrogen), and 1% (v/v) penicillin/streptomycin (Invitrogen). Human prostate cancer cell line PC3 cells, human cervical cancer cell line Hela cells, and 293T cells were obtained from the ATCC and maintained in DMEM medium supplemented with 10% (v/v) fetal bovine serum, 1% (v/v) L-glutamine, and 1% (v/v) antibiotic-antimycotic. Human non-small cell lung carcinoma cell H1299 were obtained from the ATCC and were cultured in Roswell Park Memorial Institute (RPMI) 1640 medium (Invitrogen) containing 10% (v/v) fetal bovine serum, 1% (v/v) L-glutamine, and 1% (v/v) antibiotic-antimycotic. Cells were maintained in a humidified atmosphere containing 5% CO_2_ at 37°C, and the culture medium was replaced every 2 days.

### 2.3. MTT assays and Boyden chamber assays

Cytotoxicity was determined using the 3-(4,5-dimethylthiazol-2-yl)-2,5-diphenyltetrazolium bromide (MTT; Sigma-Aldrich) assays. 3 × 10^3^ cells/well were seeded in 96-well plates. After incubation for 24 h, cells were treated with isoplumbagin at various concentrations and incubated for a further 48 h. At the end of treatment, MTT was added to each well. After incubation for 3 h, medium was removed and DMSO was added to each well to dissolve reduced MTT product formazan. The absorbance of dissolved formazan was measured at 550 nm wavelengths by a spectrophotometer. The ability of cell invasion was evaluated by Boyden chamber assay. Cells were harvested by trypsin and resuspended in a serum-free medium with 0.1% (w/v) bovine serum albumin. Then, 200 ml of cell suspension (2 × 10^5^ cells) was seeded into the upper chamber of SPLInsert with Polyethylene terephthalate (PET) membrane (pore size: 8.0 μm) (SPL Lifesciences, Korea) pre-coated with matrigel (BD Biosciences, San Jose, CA). The bottom well of 24-well plate was filled with cell medium containing 10% (v/v) fetal bovine serum with or without isoplumbagin. DMSO or isoplumbagin were separately added in the suspension of cells in the upper chamber. After incubation for 24 h, the cells migrated to the bottom side of PET membrane were fixed with 4% paraformaldehyde and stained with crystal violet. Photos of stained cells on the underside of PET membrane were taken using Zeiss Observer Z1 microscope (Zeiss, Jena, Germany). Number of cells was counted using Image J. Relative invasion ability was calculated as cell number per area

### 2.4. In vivo xenograft mice model

All procedures were performed according to approved National Tsing Hua University Institutional Animal Care and Use Committee (approval number: 10602, approval date: 2017/02/17) and National Chiao Tung University Institutional Animal Care and Use Committee (approval number NCTU-IACUC-108032, approval date: 2019/07/22) protocols. Briefly, in oral orthotopic xenograft CB17/lcr-*Prkdc^scid^*/CrlNarl mice model, 1 × 10^6^ OC3-IV2 cells were harvested and re-suspended in 100 μl PBS. The cells were injected through the oral buccal mucosa into six-week-old male CB17/lcr-*Prkdc^scid^*/CrlNarl mice (National Laboratory Animal Center, Taiwan). When the tumor size in oral buccal reached approximately 25 mm^3^ at 14 days after inoculation of the OC3-IV2 cells, mice were randomly assigned into two groups: vehicle (DMSO) control group (n = 13) and 2 mg/kg isoplumbagin group (n = 10). Treatments were given via intraperitoneal injection once every 3 days in the morning for 2 weeks. During this period, mice weights and tumor volumes (length × width^2^ / 2) were recorded every 3 days. At the end of the study (on day 18), the mice were sacrificed by carbon dioxide asphyxiation.

### 2.5. Molecular docking

To perform molecular docking of NQO1 and isoplumbagin, the crystallographic structure of NQO1 (PDB code 2F1O) and isoplumbagin (PubChem CID: 375105) was applied. The complex model of NQO1 and isoplumbagin was performed using PyRx [11]. The structural model representations and docked orientations were generated by PyMOL (DeLano Scientific) and Ligplot [12].

### 2.6. NQO1 activity assays

NQO1 activity assays kit (#ab184867) was purchased from Abcam (Cambridge, MA, USA). According to the manufacturer’s instructions, cells were lysed with 1X extraction buffer containing 1 mM phenylmethylsulfonyl fluoride for 15 min on ice, and then centrifuged at 17,000 × *g* at 4°C for 20 min. The supernatant was collected and then the protein concentration of each sample was determined by bicinchoninic acid assays (Santa Cruz Biotechnology). Diluted sample and reaction buffer were added into 96-wells plate containing menadione or isoplumbagin with cofactor NADH and water-soluble tetrazolium salt [2-(4-iodophenyl)-3-(4-nitrophenyl)-5-(2,4-disulfophenyl)-2H-tetrazolium] reagent. The absorbance of reduced water-soluble tetrazolium salt was measured at 450 nm wavelengths by a spectrophotometer.

### 2.7. Immunobloting

Cell lysates of OC3, OC3-IV2, H1299, and 293T cells were lysed by RIPA buffer (50 mM Tris, pH 7.5, 1% (v/v) Triton X-100, 150 mM NaCl, 2 mM EGTA) containing 1 mM sodium orthovanadate, 1 mM phenylmethylsulfonyl fluoride, 10 ng/ml leupeptin and 10 ng/ml aprotinin. The protein concentration of each sample was determined by the bicinchoninic acid assays, and samples were separated by SDS-PAGE, followed by immunoblotted with the indicated primary antibodies and the IRDye-conjugated secondary antibody. The immunoblots were detected using the Odyssey infrared imaging system (LI-COR Biosciences).

### 2.8. Reactive oxygen species assays

Cells were treated with DMSO or isoplumbagin for 2 h and then and harvested with trypsin. Equal numbers of cells (5 × 10^5^ cells/ml) were incubated with 5 μM of dihydroethidium (DHE; Invitrogen), a cytoplasmic superoxide indicator, for 30 min at room temperature away from light. The intracellular reactive oxygen species levels were measured by BD Accuri™ C6 flow cytometer (BD Biosciences, San Jose, CA). Histograms of 20,000 events were analyzed and DHE fluorescence was measured by filter FL-2 (585 nm). The mean fluorescence intensity was calculated by BD Accuri C6 Software.

### 2.9. TUNEL assays

Cells were treated with DMSO or isoplumbagin for 12 h. After fixed with 1% (v/v) paraformaldehyde for 15 min, cells were washed with PBS for two times and added with 70% (v/v) ethanol for 30 min on ice. After that, cells were stained by the usage of TUNEL staining kit (Abcam, #ab66108, Cambridge, MA, USA) according to the manufacturer’s instructions. Briefly, cells were incubated with DNA labeling solution for 60 min at 37°C, followed by addition of PI/RNase A solution for 30 min. TUNEL-labeled cells were evaluated by BD Accuri™ C6 flow cytometer and the data were analyzed using BD Accuri C6 Software.

### 2.10. Immunofluorescence staining

Cells were fixed by 4% (v/v) paraformaldehyde and permeabilized by 0.1% (v/v) Triton X-100. Cells were incubated in blocking buffer containing 1% bovine serum albumin. After blocking, cells were incubated overnight with indicated primary antibodies against TOM20 (a mitochondria marker), and then incubated with Alexa Fluor-conjugated secondary antibodies. Nucleus was stained with DAPI, actin was stained with phalloidin and cells were mounted using Prolong Gold reagent. Images were taken by LSM800 confocal microscopes (Zeiss).

### 2.11. High-resolution respirometry

Oxygen consumption rate in intact cells suspension was measured by Oxygraph-2k system (Oroboros Instruments). DMSO or isoplumbagin treated cells were detached from the plate by trypsinization and suspended at 5 × 10^5^ cells/ml in DMEM/KSFM medium supplemented with 10% fetal bovine serum into the O2k chambers. Oxygen consumption rate of DMSO or isoplumbagin treated cells samples were measured by adding 0.5 μM oligomycin, 0.5 μM FCCP, 1 μM rotenone and 1 μM antimycin A. For the cell permeabilization and measurement of respiration in permeabilized cells, 5□ ×□ 10^5^ cells suspended in 2□mL MiR05 buffer (110□ mM D-sucrose, 0.5□ mM EGTA, 3.0□ mM MgCl_2_, 60□ mM lactobionic acid, 10□ mM KH_2_PO_4_, 20□ mM taurine, 20□ mM HEPES, 1□ g/L bovine serum albumin, pH 7.1) were added in O2k chambers. After cells had stabilized at routine respiration, plasma membrane permeabilization was performed by adding 5μM digitonin and then O2 consumption rate was measured in response to sequential additions of 10 mM glutamate and 2 mM malate or 10 mM succinate, followed by 5 M ADP, 0.5 mM oligomycin, 0.5 μM FCCP, 1 μM rotenone, 1 μM antimycin A, 0.1□ mM TMPD, 0.4□ mM ascorbate and 5 mM NaN_3_.

### 2.12. Statistical analysis

Statistical analysis of results was carried out using the Student’s *t*-test, paired Student’s t-test or one-way ANOVA. Values reflect the mean ± SD of data obtained from two independent experiments or the mean ± S.E.M. of data obtained from at least three independent experiments. Statistical significance was defined as *P*<0.05.

## 3. Results and discussion

### 3.1. Isoplumbagin suppresses proliferation and invasion of cancer cells

In this study, isoplumbagin was chemically synthesized with 95% purity from AKos GmbH Company (Supplementary Fig. 1). The effect of isoplumbagin (Fig. 1A) on cell survival among various cancer cell lines was evaluated by MTT assays. OC3-IV2, *in vivo*-selected for highly metastasized OSCC cells [10], U87 (glioblastoma), H1299 (non-small cell lung carcinoma) and PC3 (prostate cancer) cells were treated with 0, 1, 5, 10, 25, 50 and 100 μM of isoplumbagin for 48 h. Isoplumbagin treatment inhibited cell survival of OC3-IV2, U87, H1299, and PC3 cells with IC50 5.4 μM for OC3-IV2, 2.4 μM for U87, 1.5 μM for H1299, and 6 μM for PC3 cells (Fig. 1B-E). We next examined whether isoplumbagin affects cancer cell invasion by Boyden chamber assays. As shown in Fig. 1F-J, 5 to 10 μM of isoplumbagin was sufficient to suppress the invasion of OC3-IV2, H1299, PC3 and human cervical cancer Hela cells but not U87 cells. These results reveal differential sensitivity of these cancer cells to isoplumbagin in terms of survival and invasion. U87 and H1299 cell survival are more sensitive to isoplumbagin, whereas invasion of OC3-IV2 and Hela cells is largely inhibited by isoplumbagin.

**Fig. 1.**
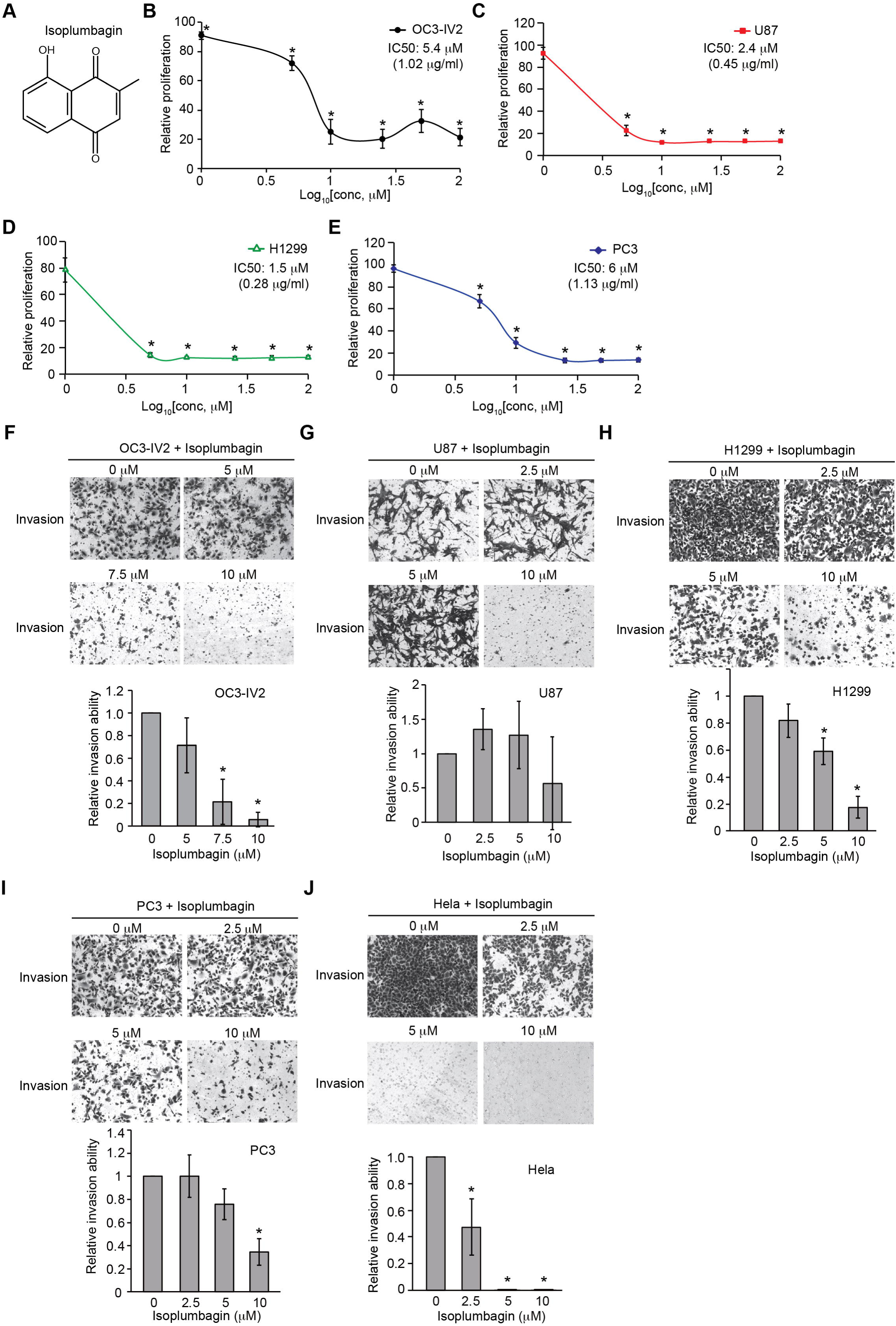
The chemical structure of isoplumbagin and its effect on survival and invasion of cancer cells. (A)Structure of isoplumbagin (5-hydroxy-3-methyl-1,4-naphthoquinone). (B-E) The MTT assay was used to determine proliferation of OC3-IV2 (B), U87 (C), H1299 (D) and PC3 (E) cells treated with isoplumbagin for 48 h. Calculated IC50 values of isoplumbagin for OC3-IV2, U87, H1299 and PC3 cells. (F-J) Invasion of OC3-IV2 (F), U87 (G), H1299 (H), PC3 (I) and Hela (J) cells treated with isoplumbagin was assessed with the Boyden chamber assays. Values for invasion were normalized to those for cells treated with solvent control (DMSO) *Compared with solvent control (DMSO). Data from three independent experiments are presented as mean ± S.E.M. (*P<0.05, paired Student’s *t*-test).

### 3.2. Isoplumbagin inhibits tumor growth in an OSCC xenograft model

To test the anti-cancer effect of isoplumbagin *in vivo*, we established orthotopic xenograft model by injecting OC3-IV2 cells to oral buccal of CB17/lcr-*Prkdc^scid^*/CrlNarl mice. When the tumor volume in oral buccal reached up to 25 mm^3^, isoplumbagin was intraperitoneally injected once every three days for 2 weeks. The body weight of isoplumbagin- and vehicle-treated mice was similar (Fig. 2A), indicative of no obvious systemic toxicity at this treatment regimen. On the other hand, the average tumor weights for mice treated with 2 mg/kg of isoplumbagin were reduced (Fig. 2B-C) and the tumor sizes in mice treated with isoplumbagin were approximately 50% smaller compared to control mice. During the course of 18-day treatment regimen, the average tumor volume in 10 mice treated with 2 mg/kg of isoplumbagin groups was decreased compared to the vehicle-treated 11 mice (Fig. 2D). Due to the difficulty of measuring tumor volume in the small oral cavity of mice, the variation of tumor volume is relatively large rendering no statistical difference. Nonetheless, these *in vivo* data indicate that 2 mg/kg isoplumbagin inhibits the growth of OSCC-derived tumors without systemic toxicity.

**Fig. 2.**
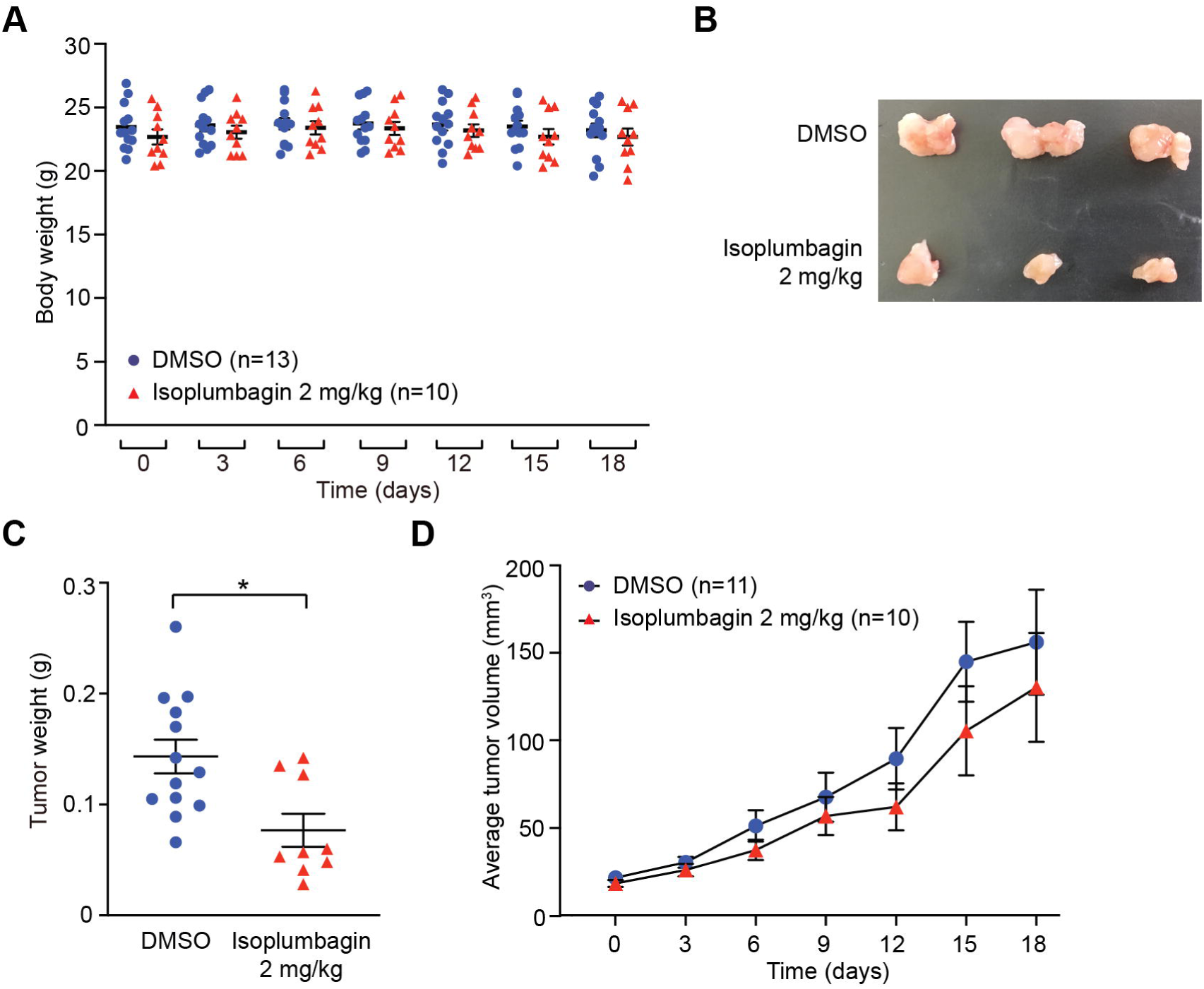
The effect of isoplumbagin in orthotopic xenograft mice model. (A) OC3-IV2 cells were injected into the oral buccal mucosa of CB17/lcr-*Prkdc^scid^*/CrlNarl mice. After tumor volume reached approximately 25 mm^3^, mice were treated with vehicle (DMSO, n = 13) or isoplumbagin (2 mg/kg, n = 10) via intraperitoneal injections once every 3 days. Body weight of mice was measured from day 0-18 after treatment. Each value in the graph represents the mean ± S.E.M from mice. (B-C) Mice were sacrificed and tumors were isolated from the vehicle (DMSO, n = 13) or isoplumbagin treatment (n = 9) at 18 days. Representative photograph of tumors was shown in (B). The tumor weight (C) of harvested tumor was measured from (B). Each value in the graph represents the mean ± S.E.M from harvested tumor (\**P*<0.05, Student’s t-test). (D) Tumor volume from (A) with vehicle-(DMSO, n = 11) or isoplumbagin-treated (n = 10) mice were measured during the course of 18 days treatment. Statistical analysis of results in (D) was carried out using the one-way ANOVA (Tukey’s Test). Each value in the graph represents the mean ± S.E.M.

### 3.3. Isoplumbagin is a substrate of NAD(P)H dehydrogenase [quinone] 1 (NQO1)

To identify the candidate targets for isoplumbagin, we used prediction softwares, Swiss Target Prediction, Pharmmapper, Polypharmacology Browser and Similarity ensemble approach, followed by the Database for Annotation, Visualization and Integrated Discovery (DAVID) to investigate of the interaction network (Fig. 3A). The top five functions of isoplumbagin-targeted proteins were related to oxidation-reduction process, signal transduction, response to drug, negative regulation of apoptosis process and protein phosphorylation (Fig. 3B). Based on the cluster and the highest enrichment score, NQO1 protein was ultimately chosen for further investigation. Physically, NQO1 enzyme functions as a homodimer [13, 14] that uses the NADH or NADPH as a cofactor to catalyze the reduction of quinones to hydroquinones. To understand how isoplumbagin interacts with NQO1 homodimer, the molecular docking and ligand binding simulation approach was used to examine their probable interaction. Four combination of NQO1 homodimer are shown based on eight chains of crystal structure of NQO1 (PDB code 2F1O). Isoplumbagin bound to the enzymatic active site of NQO1 homodimers through hydrogen bonds with residues Tyr126, Tyr128, or His161 and van der Waals interactions with other residues Trp105, Phe106, Met131, Gly149, Gly150, Met154, His161, Phe178, and Phe236 of NQO1 (Fig. 3C). Based on their predicted binding affinity and the distance of hydrogen bonds, we propose the most stable complex is isoplumbagin and B/D chains of NQO1 (Fig. 3D). Compare to the previously reported interaction between NQO1 and its inhibitor dicoumarol, residues Trp105, Tyr126, Tyr128, Gly149, His161 and Phe178 of NQO1 were involved [15]. We next investigated whether isoplumbagin is a substrate or an inhibitor of NQO1 using an *in vitro* NQO1 activity assays. This assay uses menadione as a substrate of NQO1, together with cofactor NADH, and reduces menadione for further reduction of water-soluble tetrazolium salt to yellow color formazan. As shown in Fig. 3D, formazan was increased in the presence of menadione, compared to no substrate, similar to the effect of isoplumbagin. This result suggests that isoplumbagin is a NQO1 substrate.

**Fig. 3.**
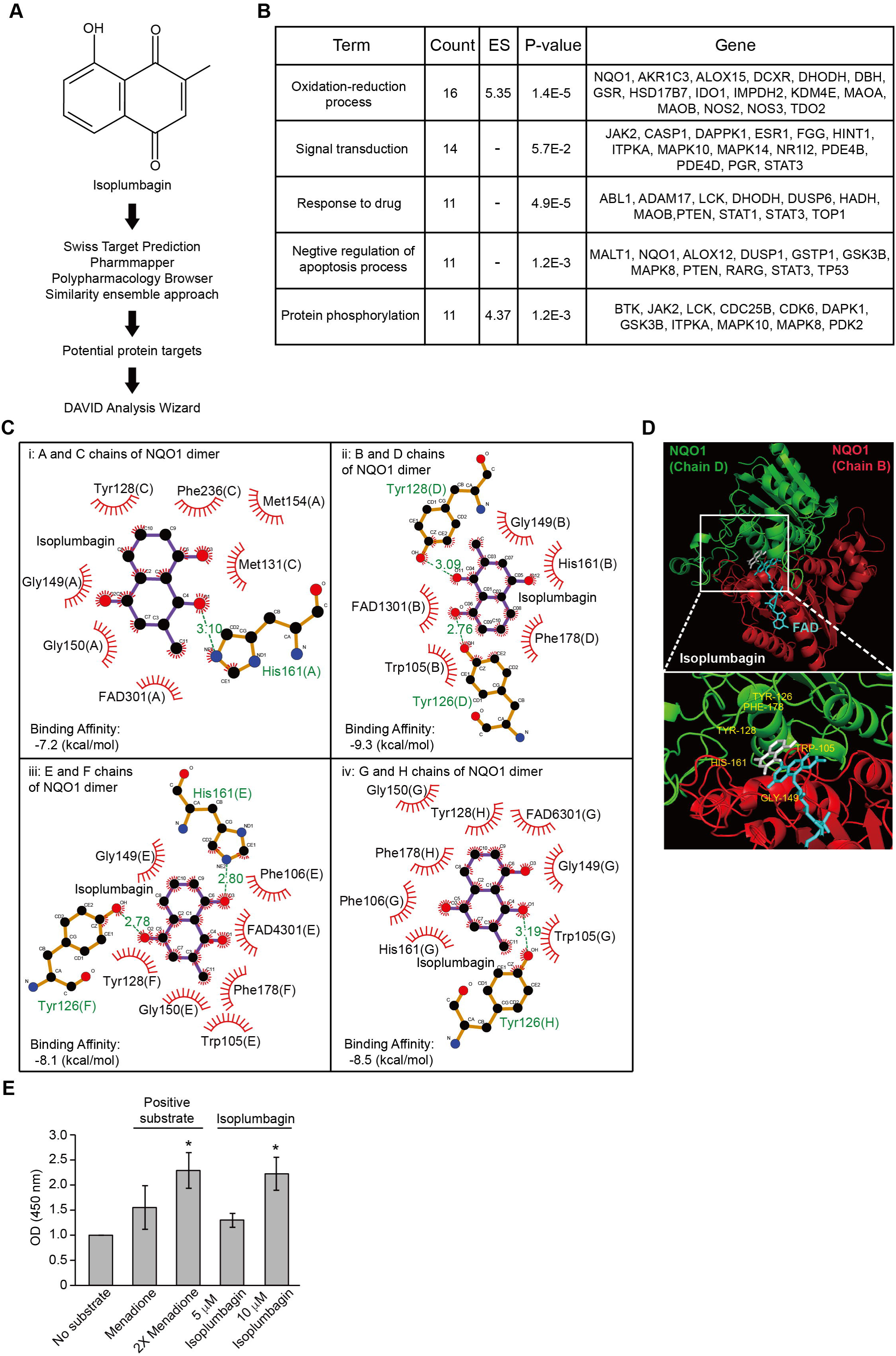
Isoplumbagin is a substrate of NQO1. (A) The target prediction of isoplumbagin used target prediction softwares, Swiss Target Prediction, Pharmmapper, Polypharmacology Browser and Similarity ensemble approach, followed by the Database for Annotation, Visualization and Integrated Discovery (DAVID). (B) The Gene Ontology (GO) term enrichment results of potential binding targets for isoplumbagin based on DAVID analysis. (C) Schematic representation of isoplumbagin and NQO1 homodimer interaction: i for A and C chains, ii for B and D chains, iii for E and F chains, and iv for G and H chains of NQO1 homodimers. FAD is cofactor of NQO1. Calculated binding affinity of isoplumbagin and NQO1 homodimer was shown. Hydrogen bonds and van der Waals interactions are shown in stick representation and decorated arc, respectively. (D) The molecular docking between isoplumbagin and B/D chains of NQO1. Isoplumbagin is colored in white; NQO1 homodimer is colored in red for chain B and green for chain D; FAD is in blue. (E) The cytosolic fractions of OC3-IV2 cells were used for the NQO1 enzymatic activity assay. The NQO1 enzyme activity was calculated by measuring the simultaneous reduction of menadione or isoplumbagin with cofactor NADH and water-soluble tetrazolium salt (a highly sensitive tetrazolium reagent) which leads to increased absorbance at 450 nm. *Compared with control (no substrate). Data from three independent experiments are presented as mean ± S.E.M. (*P<0.05, paired Student’s *t*-test).

### 3.4. NQO1 levels varies in different types of cancer

NQO1 facilitates detoxification of quinones to stable hydroquinones through two-electron reduction process (Supplementary Fig. 2A). The two-electron reduction catalysed by NQO1 protects cells from oxidative stress thereby avoiding the production of semiquinone radicals. This mechanism has been used in a number of cancer cell types, such as lung, pancreatic and breast cancer, for survival [16, 17]. Interesting, we found that NQO1 was increased in OC3 cells and H1299 compared to normal 293T cells, but NQO1 expression was significantly reduced in highly invasive OC3-IV2 cells compared with its parental OC3 cells (Fig. 4A). In addition, analysis of public Gene Expression Omnibus (GEO) datasets for NQO1 expression in patient samples of various cancer types showed different expression profiles in primary tumors compared with their respective normal tissues or metastatic stage; NQO1 expression was decreased in colorectal cancer and nasopharyngeal carcinoma, increased in pancreatic tumor and metastatic OSCC, and no different in tongue squamous cell carcinoma, papillary thyroid cancer and metastatic stage of cervical cancer and melanoma (Fig. 4B-C). These results suggest that high or low NQO1 might not be a bonafide prognostic marker. Nonetheless, lower NQO1 expression correlates with advanced prostate cancer, metastatic tissues and mesenchymal attributes [18]. Thus, targeting NQO1 should carefully consider the stage/progress of the diseases.

**Fig. 4.**
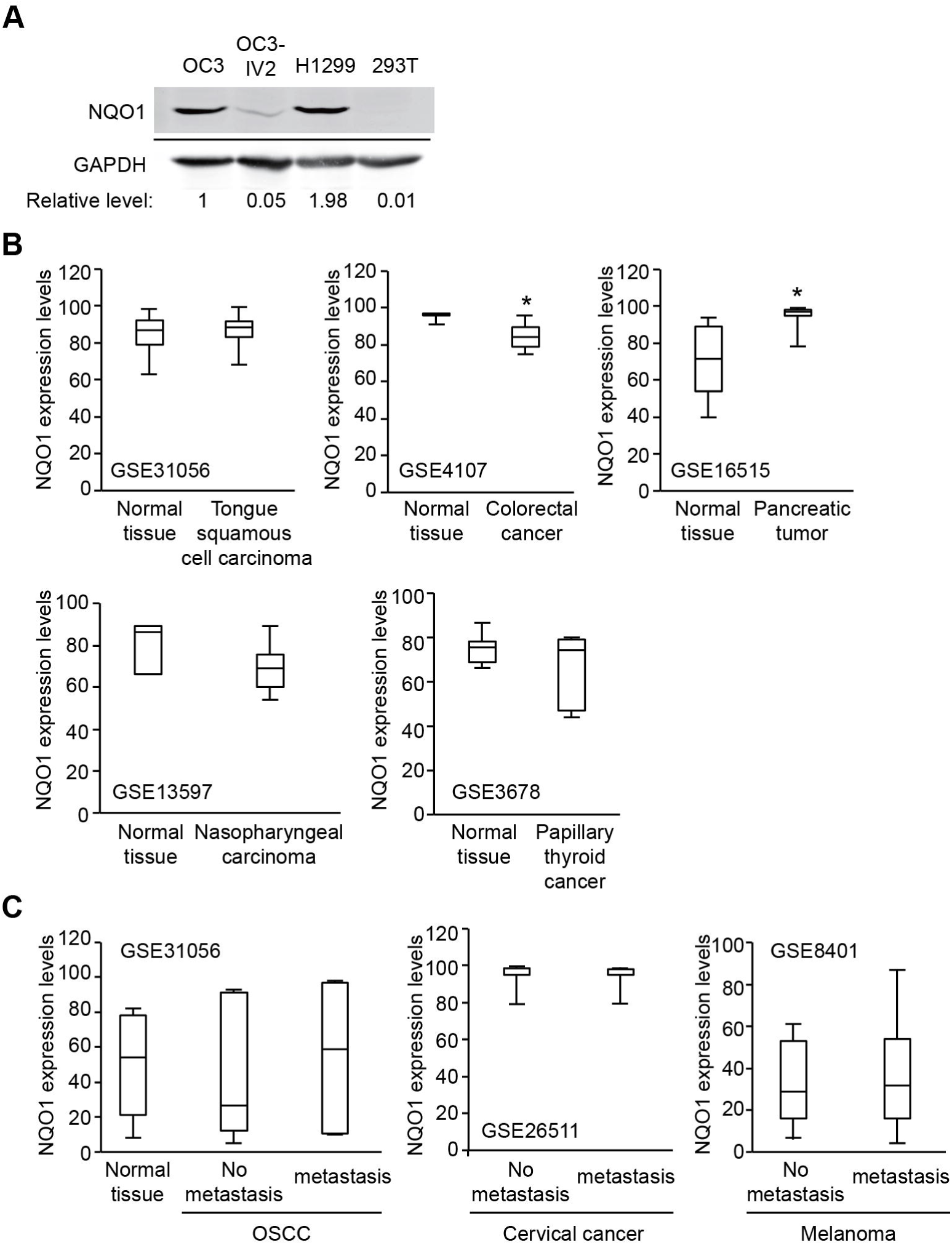
NQO1 levels in cancer cell lines and in patient tissues. (A) Cell lysates of OC3, OC3-IV2, H1299 and 293T cells were resolved via SDS-PAGE followed by immunoblotting with anti-NQO1 and anti-GAPDH antibodies. GAPDH was used as a loading control. Relative level of NQO1 was normalized to GAPDH level. (B) Gene expression data obtained from Gene Expression Omnibus (GEO) database were used to analyze NQO1 expression in tumor and normal tissues. Tongue squamous cell carcinoma: GSE31056, colorectal cancer: GSE4107, pancreatic tumor: GSE16515, nasopharyngeal carcinoma: GSE13597, papillary hyroid cancer: GSE3678. (C) Gene expression of NQO1 in tumor and metastasis from GEO database. OSCC: GSE31056, cervical cancer: GSE26511, melanoma: GSE8401. *Compared with respective normal tissues (*P<0.05, Student’s *t*-test).

Dicoumarol, an available NQO1 inhibitor, which competes with NAD(P)H for binding to NQO1, had no obvious effect on suppression of OSCC growth compared to isoplumbagin (Supplementary Fig. 3A and Fig. 1B). This phenomenon implies that inhibition of NQO1 by dicoumarol is not a useful approach for treatment of highly invasive OSCC, which also suggests that isoplumbagin may serve as a bioactivator/substrate for NQO1 rather than as an inhibitor.

### 3.5. Isoplumbagin does not increase oxidative stress nor DNA fragmentation in the highly invasive oral cancer cells

What is the molecular mechanism of isoplumbagin-induced NQO1 action on suppression of cancer cell growth and invasion? β-lapachone, which is currently under multiple Phase I/II clinical trials, is known to be bioactivated by NQO1 in NQO1^high^ tumors leading to generation of reactive oxygen species [19–21]. Elevated oxidative stress is correlated with tumor initiation, metastasis and therapeutic resistance. Hence, we examined whether a decrease in cell viability of OC3-IV2 cells treated with isoplumbagin was due to the NQO1-induced intracellular oxidative stress accumulation. As shown in Fig. 5A, OC3-IV2 cells showed no accumulation of reactive oxygen species with isoplumbagin treatment. Another mechanism by which unstable hydroquinone exerts cytotoxicity may be as alkylating agents like mitomycin C to induce DNA crosslinking leading to DNA damage. We, therefore, assessed the effect of isoplumbagin on DNA damage in OSCC cells by TUNEL assays. Our results also showed no induction of DNA fragmentation by isoplumbagin in OC3-IV2 cells (Fig. 5B). Considering the lower level of NQO1 in OC3-IV2 cells compared to OC3 cells, these data are not surprising. We think isoplumbagin may exert anti-cancer effect in the highly invasive OSCC via a novel mechanism.

**Fig. 5.**
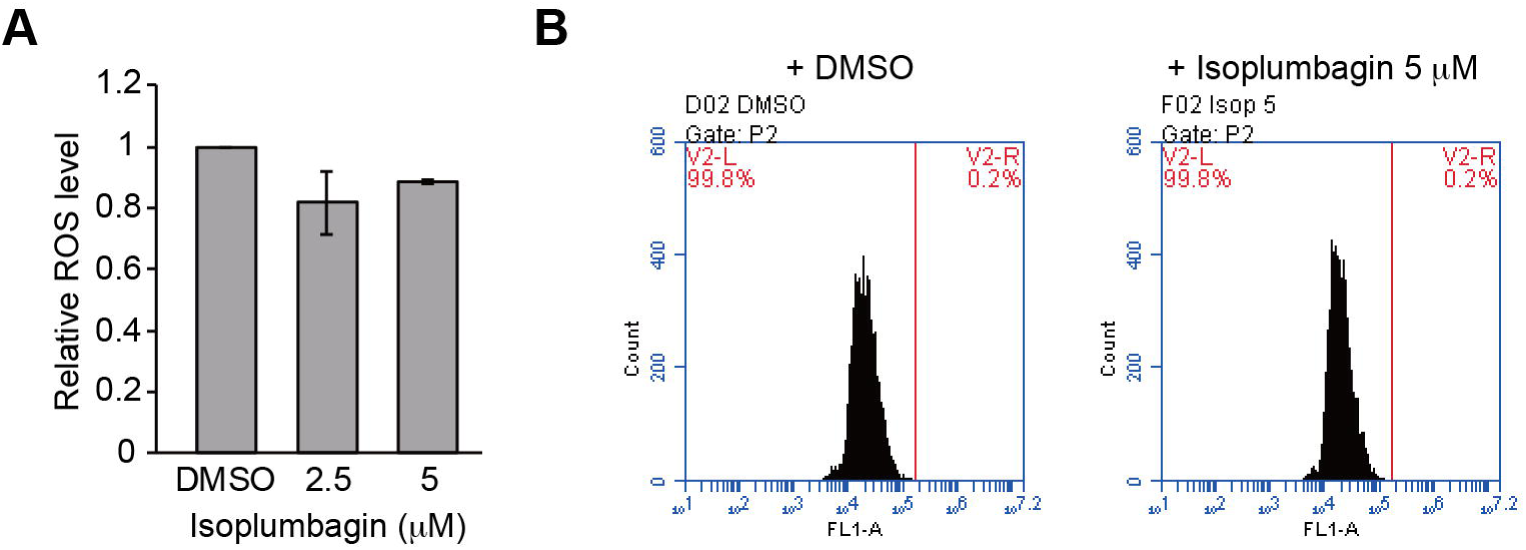
The effect of isoplumbagin on intracellular oxidative stress and DNA damage of OSCC cells. (A) OC3-IV2 cells were treated with either solvent control (DMSO) or isoplumbagin for 2 h. The intracellular reactive oxygen species levels were determined using the fluorescent dye DHE via flow cytometry. The mean fluorescence intensity were normalized to the DHE fluorescence in the DMSO control group. Data from two independent experiments are presented as mean ± SD. (B) OC3-IV2 cells were treated with either DMSO or isoplumbagin for 12 h. The DNA fragmentation was determined by TUNEL analysis. Representative data were from two independent experiments.

### 3.6. Isoplumbagin regulates mitochondrial morphogenesis and respiration in highly invasive OC3-IV2 cells

We previously reported that OC3-IV2 cells expressed high level of ROS1 oncogene compared to OC3 cells [10]. The high level of ROS1 oncoprotein did not increase oxidative stress. Our recent findings revealed that ROS1 localized to the mitochondria and increased mitochondrial fission [22]. We thus investigated whether the effect of isoplumbagin on invasive OSCC is associated with mitochondrial dysfunction. First, the effect of isoplumbagin on mitochondrial morphogenesis of OC3-IV2 cells was examined by immunofluorescence staining. In the isoplumbagin-treated OC3-IV2 cells, the mitochondria became elongated (presumably through increasing fusion), compared to the mostly fragmented phenotype in mock-treated cells (Fig. 6). Since mitochondrial fragmentation is often associated with cancer metastasis [23] and stemness [24, 25], isoplumbagin may target regulators of mitochondrial morphogenesis to exert anti-invasiveness and anti-stemness properties.

**Fig. 6.**
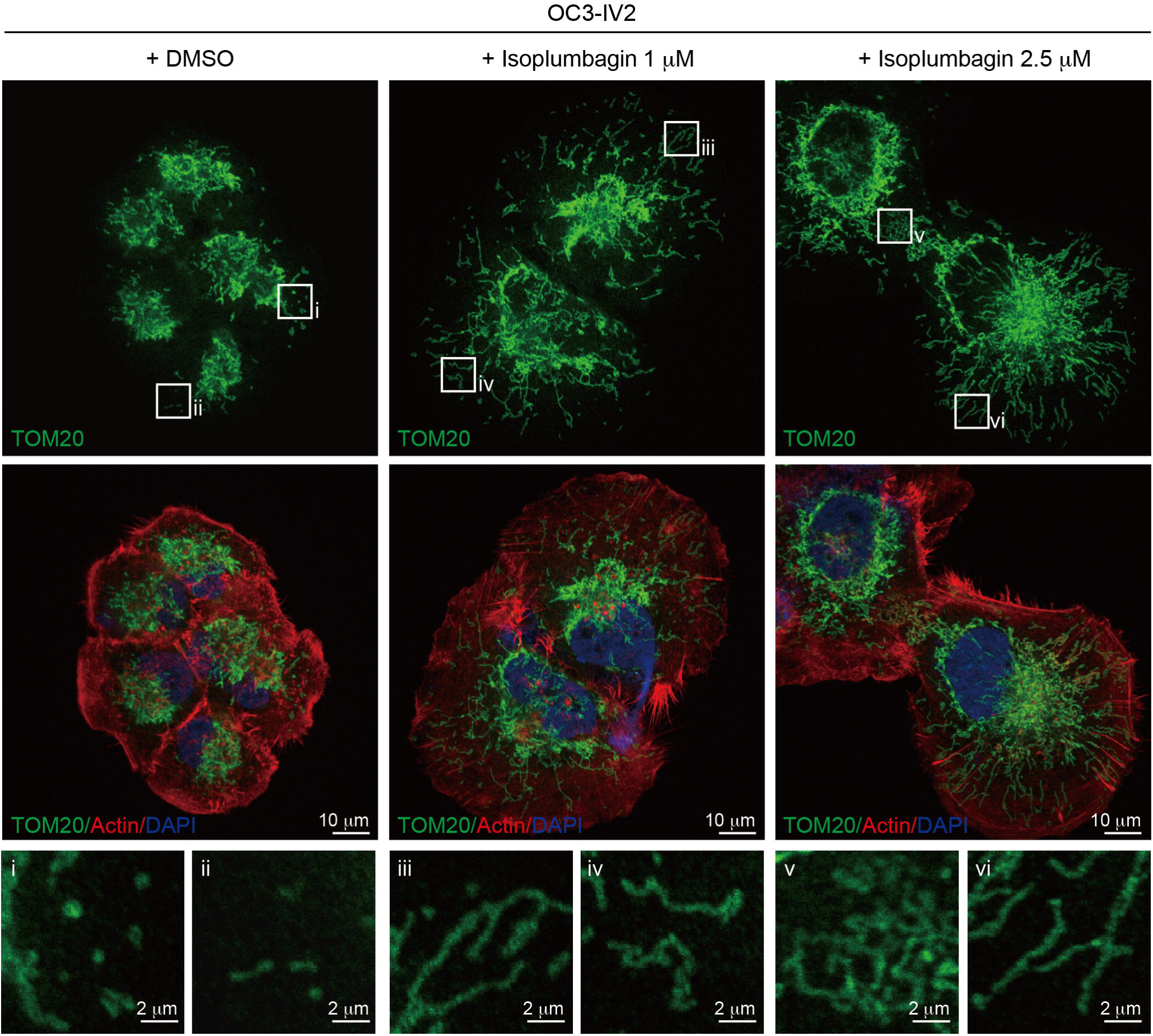
The effect of isoplumbagin on mitochondrial morphogenesis of OC3-IV2 cells. Mitochondrial morphologies of OC3-IV2 cells treated with either DMSO or isoplumbagin were determined by immunofluorescence staining using anti-TOM20 (mitochondria, green), phalloidin (actin, red), and DAPI (nucleus, blue). Enlarged panels from the boxed area are shown in the bottom panel. Images were taken using Zeiss LSM800. Scale bar: 10 μm; 2 μm for enlarged images.

We next examined the intact cell oxygen consumption rate (OCR) of DMSO-treated or isoplumbagin-treated OC3-IV2 cells by high-resolution respirometry using an Oxygraph-2k to determine whether isoplumbagin influenced mitochondrial respiration (OXPHOS). As shown in Fig. 7A, isoplumbagin decreased basal mitochondrial OXPHOS activity. In addition, inhibiting ATP synthase activity by oligomycin was associated with a decrease of ATP generation by 24% in isoplumbagin-treated OC3-IV2 cells. Subsequently, addition of the uncoupler FCCP revealing the maximal capacity of the electron transport system (ETS) inhibited 38% after isoplumbagin treatment, whereas spare respiratory capacity, defined as the difference between basal respiration and FCCP-induced maximal respiration, was reduced 44% by isoplumbagin. Finally, rotenone and antimycin A fully blocked respiration through inhibiting Complexes I and III. These data indicate that isoplumbagin suppresses basal respiration, maximal respiration, spare respiratory capacity, and ATP production.

**Fig. 7.**
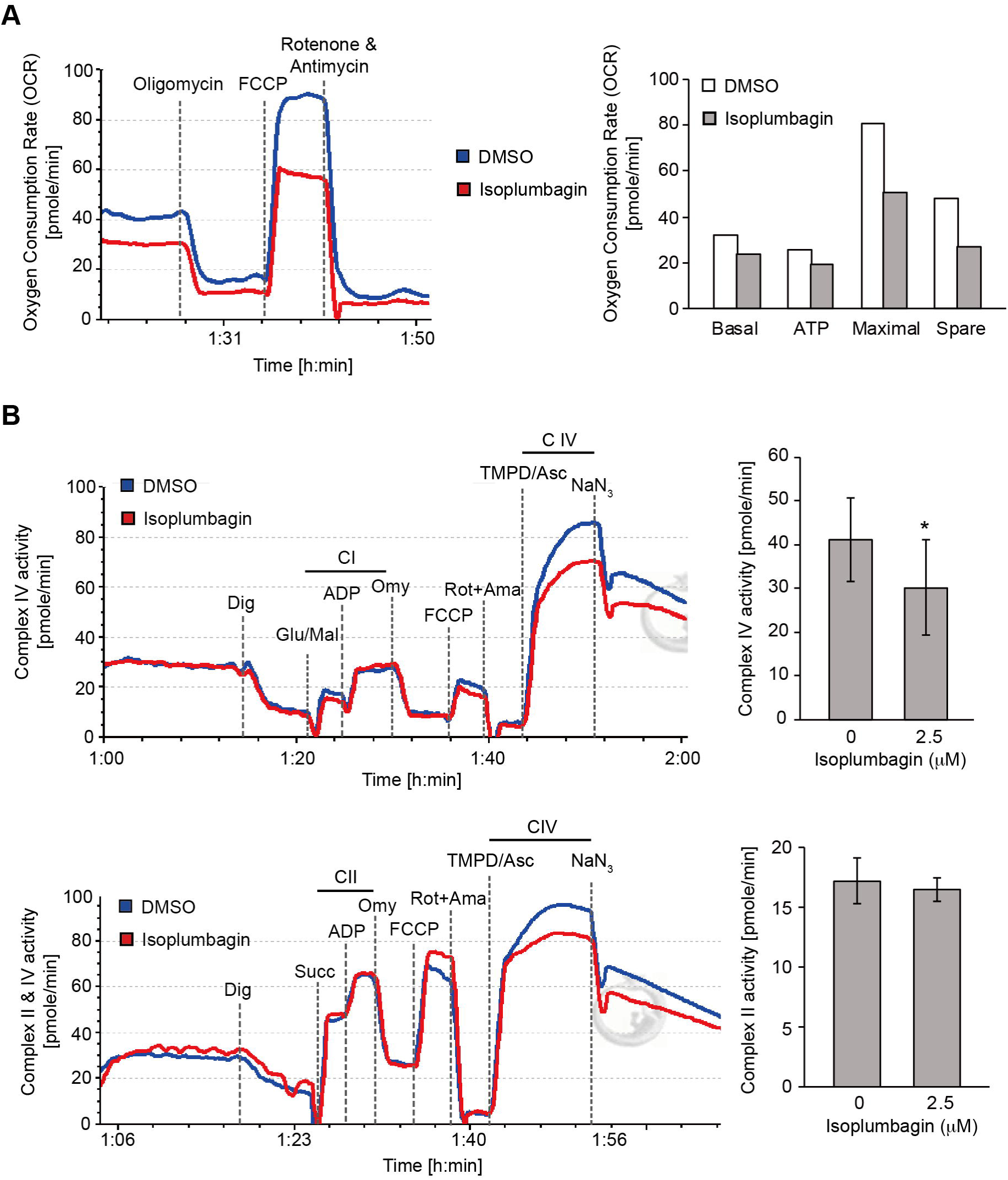
The effect of isoplumbagin on mitochondrial respiration and complex activity of OC3-IV2 cells. (A) OCR of OC3-IV2 cells treated with DMSO (as control) or 2.5 μM isoplumbagin were measured by Oroboros instruments. This result was from one independent experiment. (B) OC3-IV2 cells treated with DMSO or 2.5 μM compound isoplumbagin were permeabilized with digitonin (Dig) and added indicated substrates and inhibitors, and then mitochondria complex I/IV (upper panel) or II/IV (bottom panel) activity were determined by Oroboros instruments. Glu/Mal: glutamate and malate, succinate, ADP, Omy: oligomycin, FCCP, Rot: rotenone, Ama: antimycin A, TMPD: tetramethyl-p-phenylenediamine, Asc: ascorbate and NaN_3_. Data from three independent experiments are presented as mean ± S.E.M. (*P<0.05, paired Student’s *t*-test)

To directly examine how isoplumbagin reduced mitochondrial OXPHOS, the individual OXPHOS complexes activity following a substrate uncoupler inhibitor titration (SUIT) protocol [26] that was performed to permeabilized OC3-IV2 cells with digitonin and measured by high-resolution respirometry. The glutamate/malate-stimulated complex I linked respiration (Fig. 7B, upper panel) and succinate-supported complex II linked respiration (Fig. 7B, bottom panel) were not altered by isoplumbagin treatment, but TMPD/ascorbate-stimulated respiration of complex IV was decreased in isoplumbagin-treated OC3-IV2 cells compared to DMSO-treated cells. Taken together, these data suggest that isoplumbagin reduces mitochondrial OXPHOS through inhibiting complex IV activity. These data are consistent with our findings that the highly invasive OSCC correlated with enhanced mitochondrial fission, biogenesis and cell metabolic plasticity [22].

## 4. Conclusion

These findings suggest that isoplumbagin is a potential anti-cancer chemical against human oral squamous cell carcinoma, glioblastoma, non-small cell lung carcinoma, prostate and cervical cancers. We identify a novel mechanism of isoplumbagin effect through acting as a NQO1 substrate to modulate mitochondrial morphogenesis and respiration in highly invasive OSCC cells.

## Acknowledgements

We thank Professor Shih-Che Sue and Dr. Yi-Zong Lee at the National Tsing Hua for validating isoplumbagin compound using high performance liquid chromatography analysis. We also thank Dr. Cecilia Koo Botanic Conservation Center in Taiwan for providing reagents.

---

**Supplementary Fig. 1.**
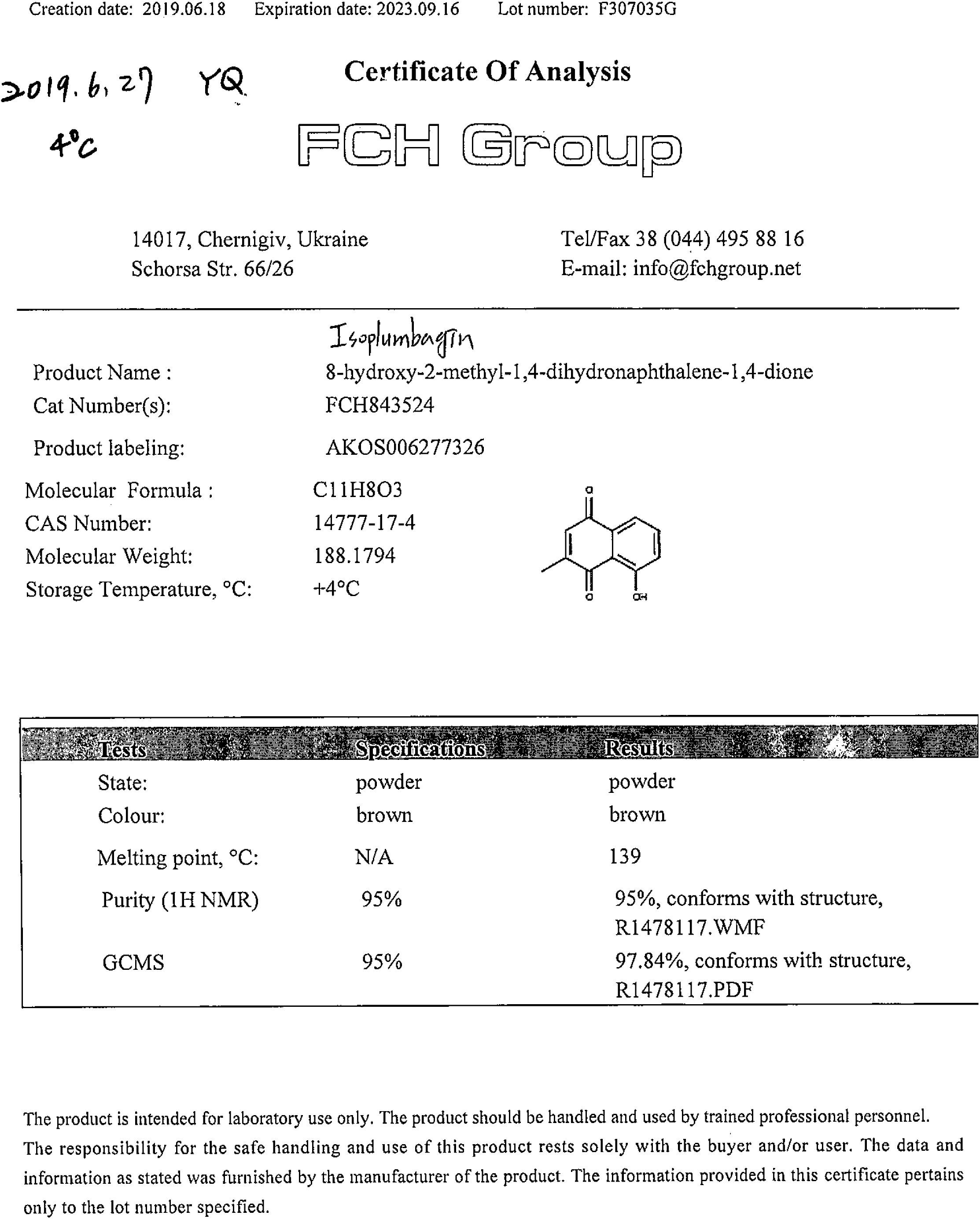
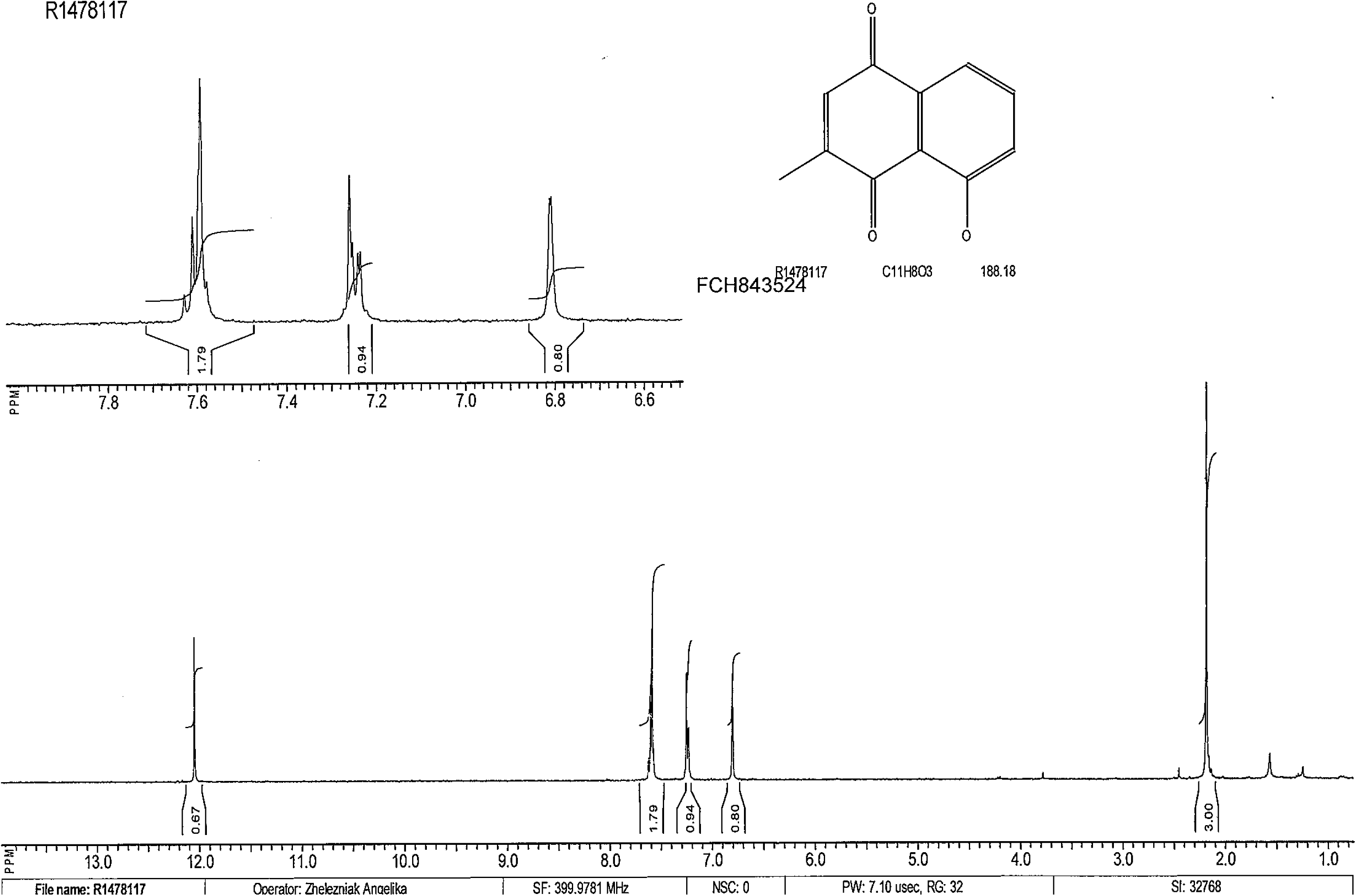
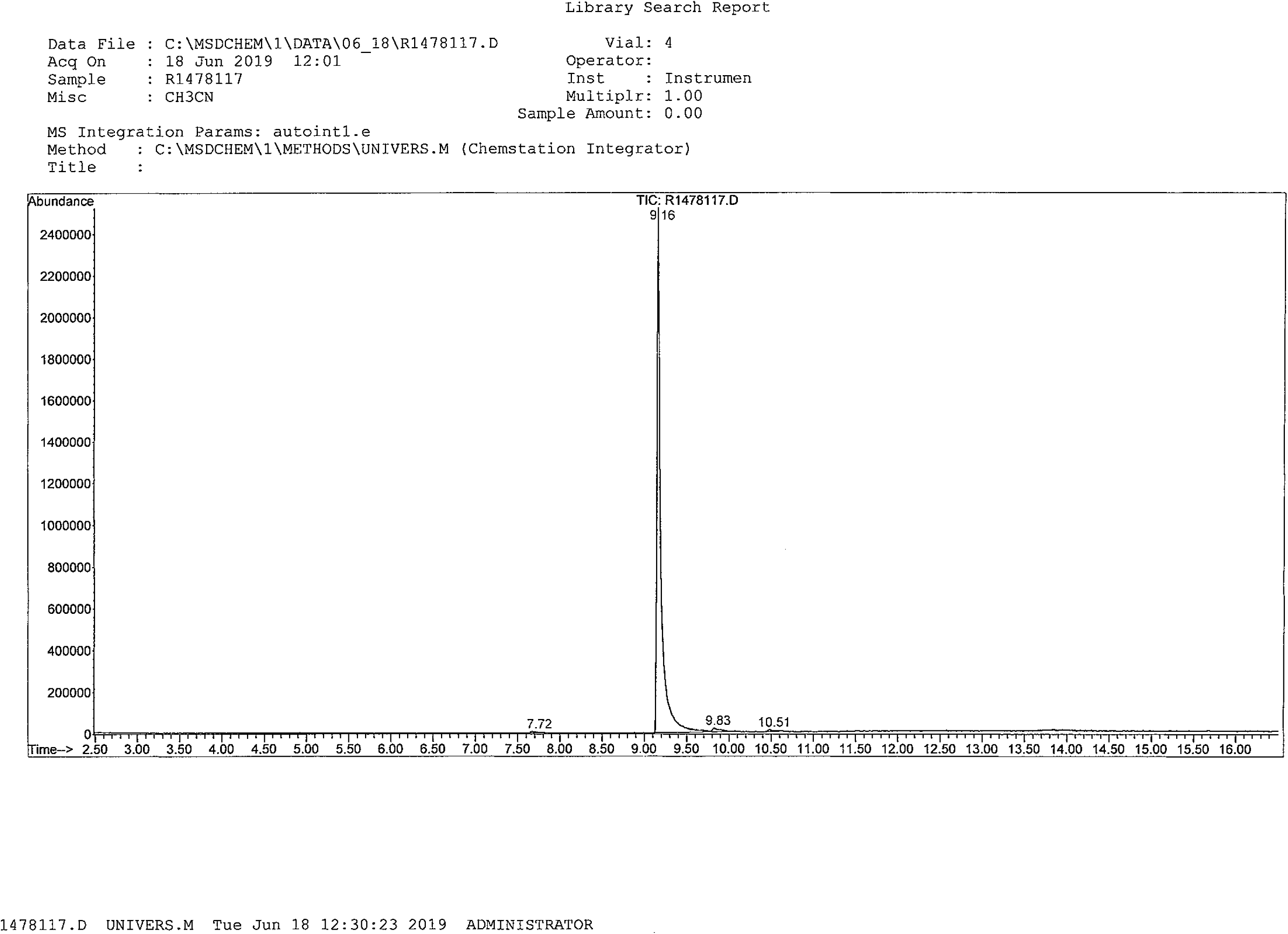
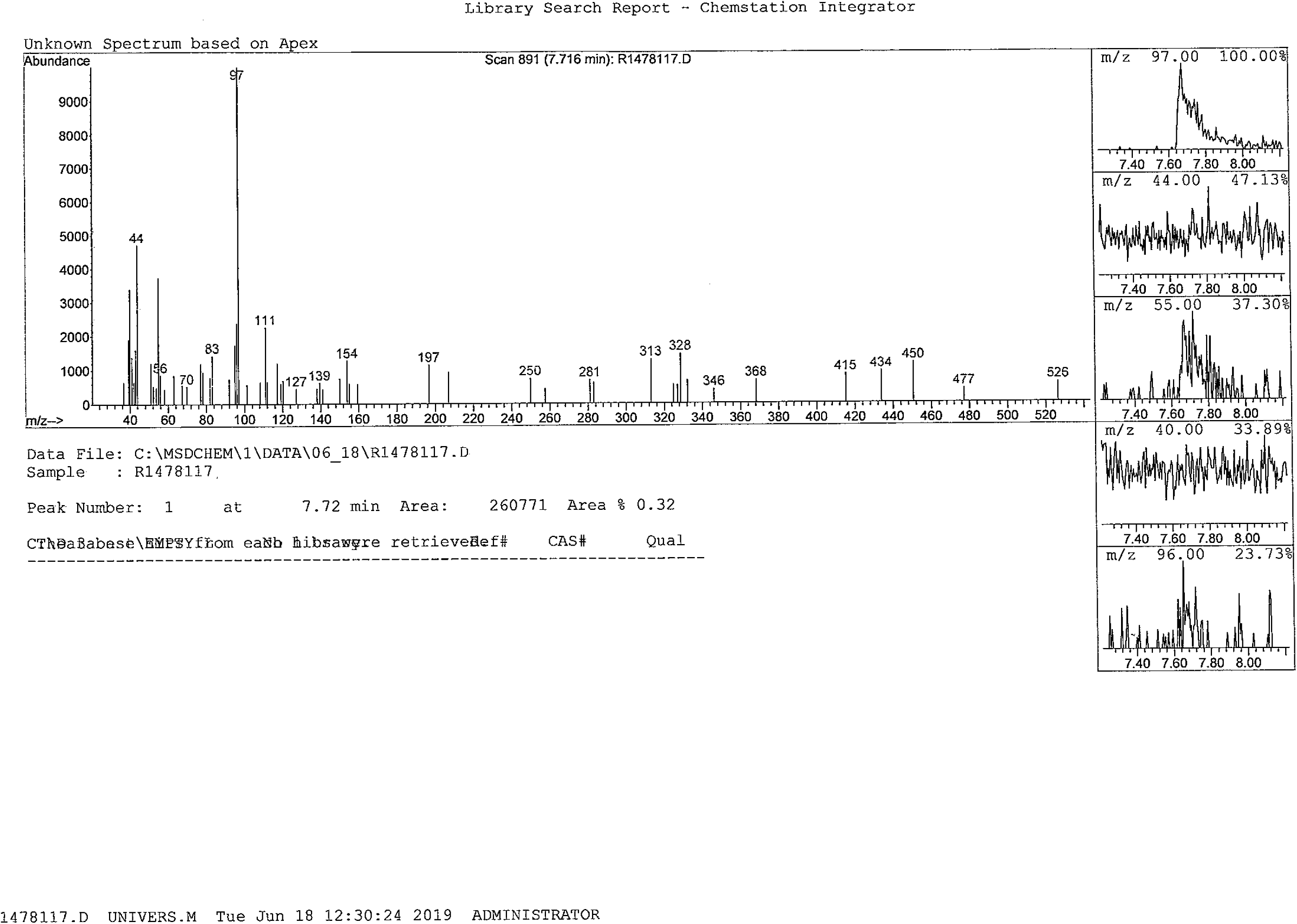
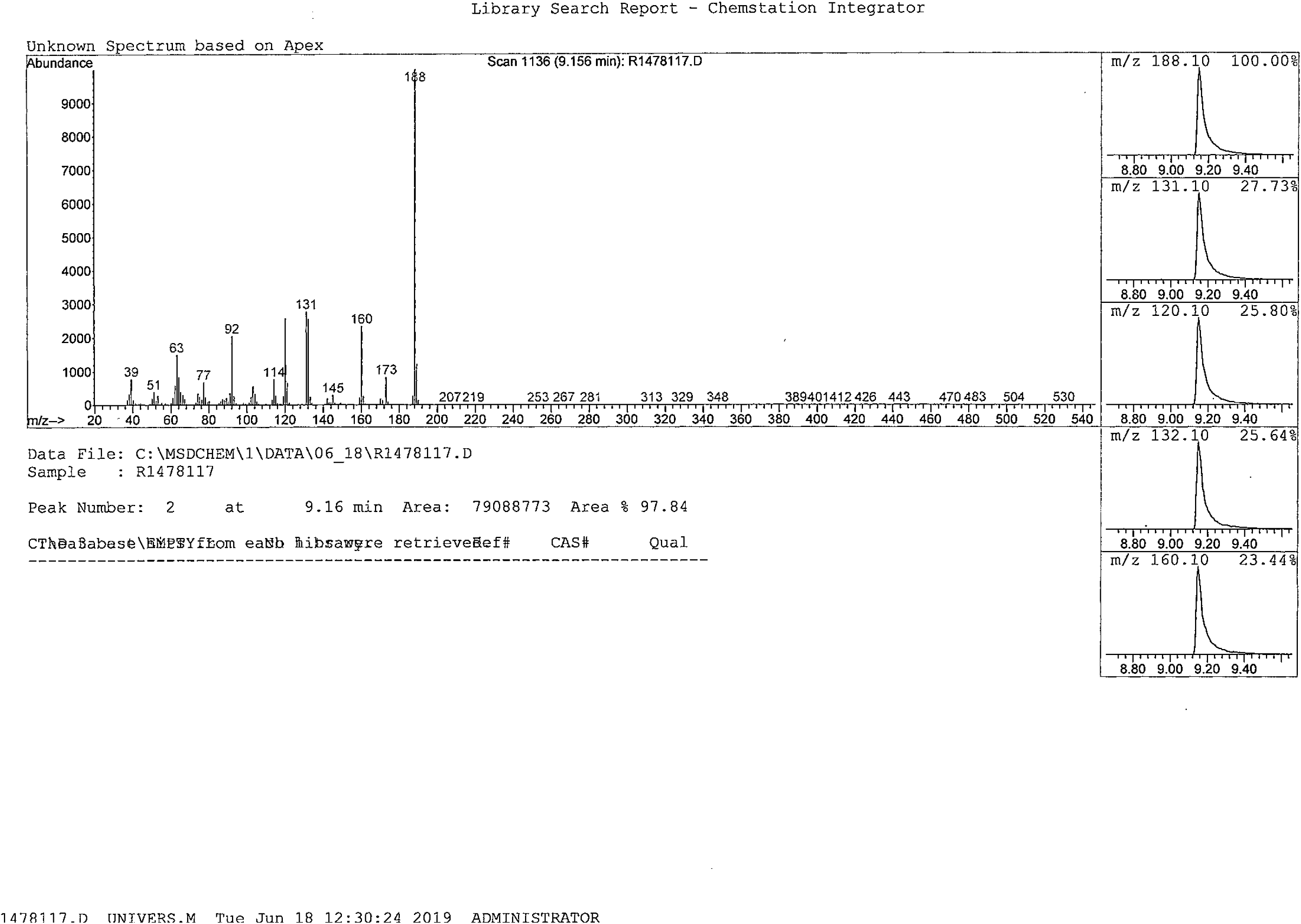
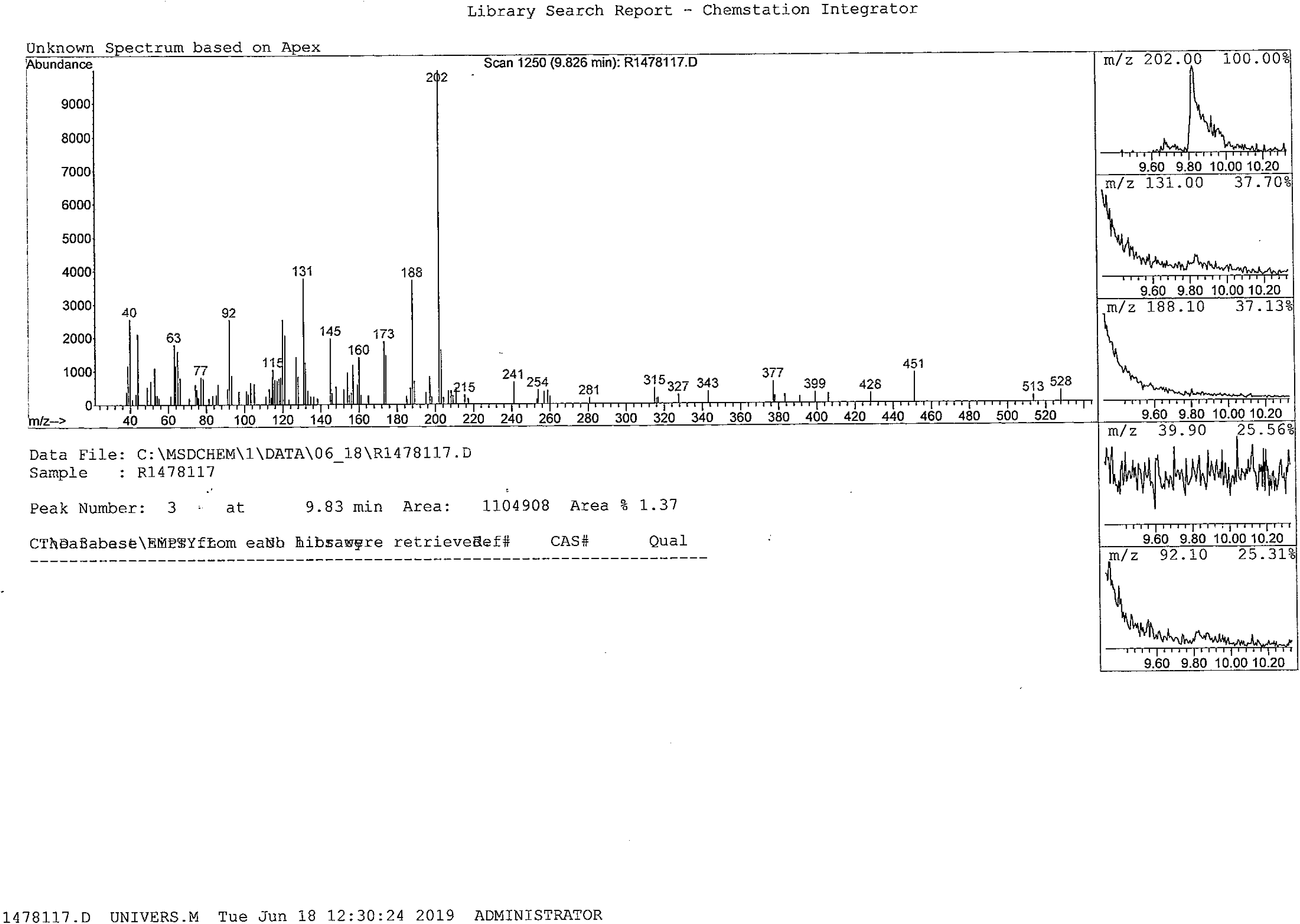
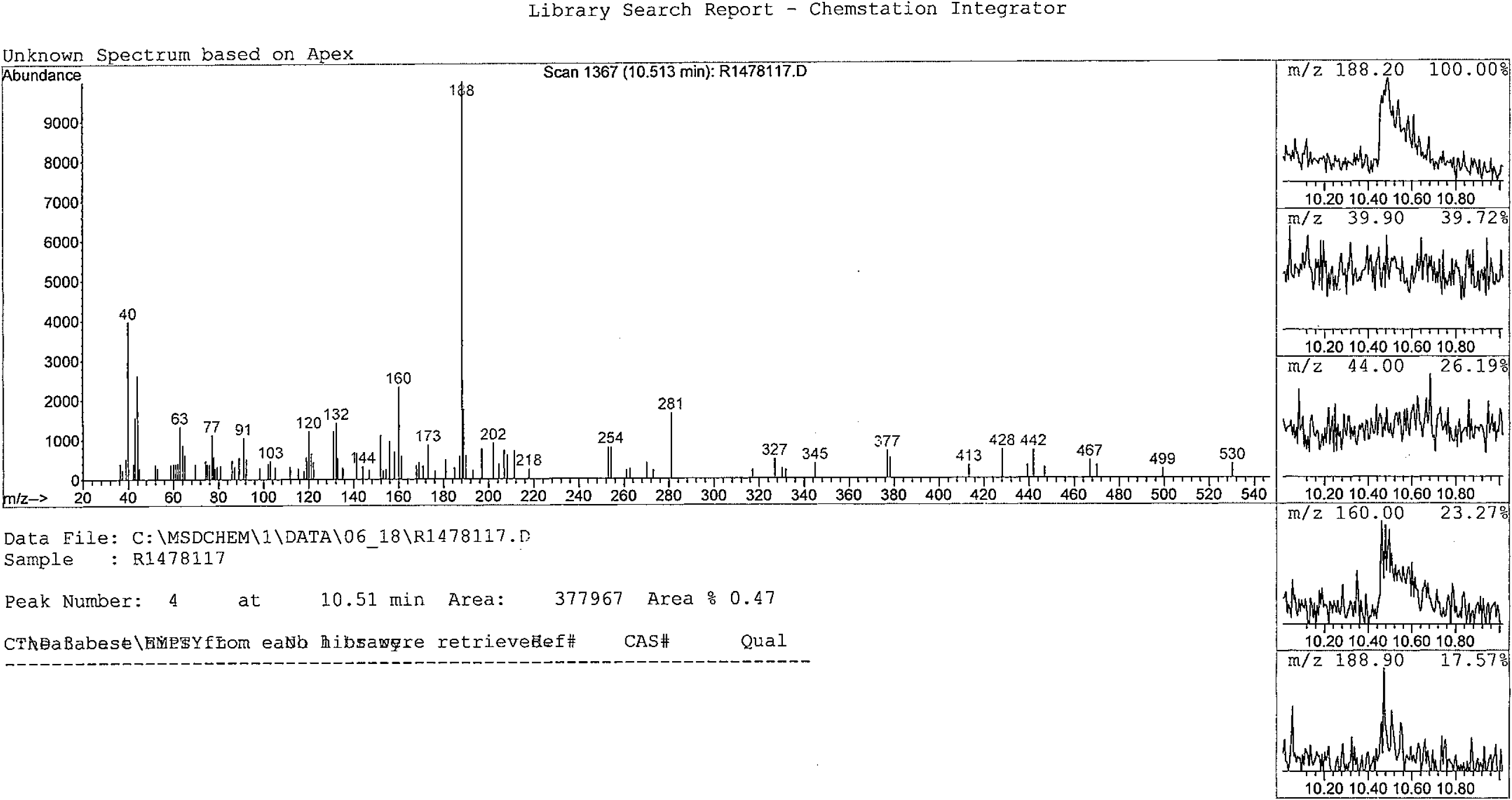
The chemically synthesized isoplumbagin and HPLC analysis.

**Supplementary Fig. 2.**
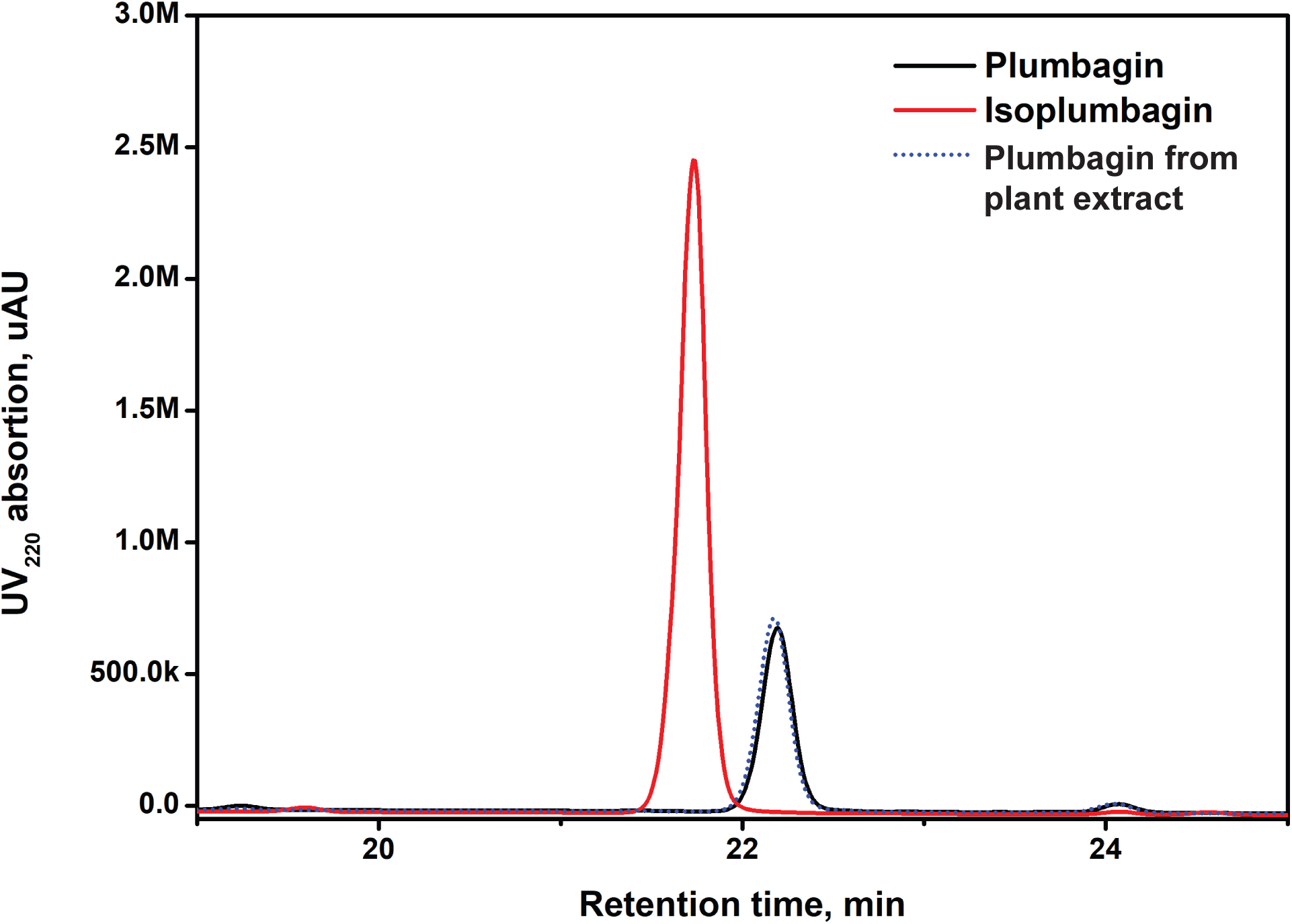
Scheme of two electron reduction of isoplumbagin by NQO1 generates the hydroquinone.

**Supplementary Fig. 3.**
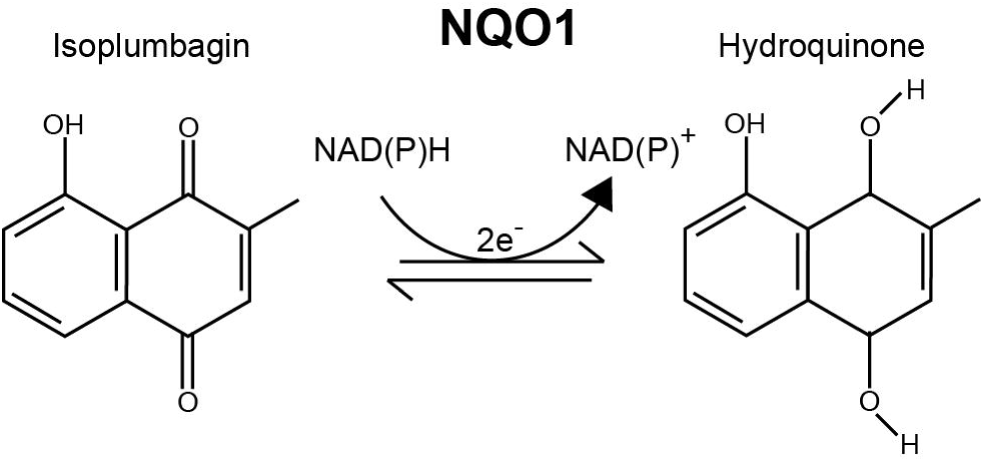
The relative NQO1 expression and the effect of dicumarol on survival of OSCC cells. Proliferation of OC3-IV2 cells treated with dicoumarol for 48 h was measured with the MTT assay. Data from three independent experiments are presented as mean ± S.E.M.

**Supplementary Fig. 4.**
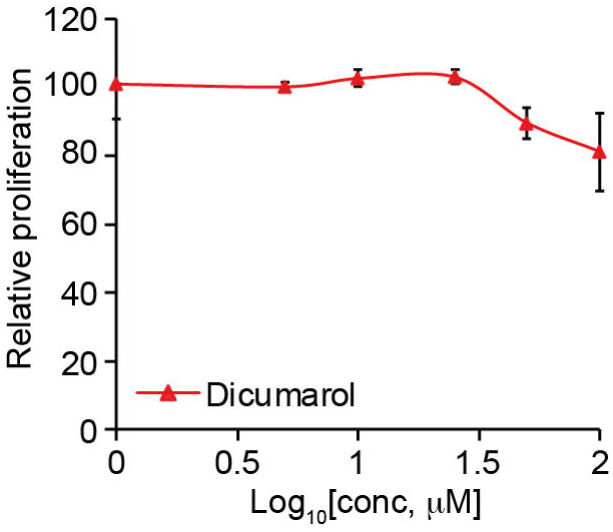
Uncropped images of the original western blots for Fig. 4A.

